# CRISPR-based dissection of miRNA binding sites using isogenic cell lines is hampered by pervasive noise

**DOI:** 10.1101/2024.09.03.611048

**Authors:** Mahendra K. Prajapat, Joana A. Vidigal

## Abstract

Non-coding regulatory sequences play essential roles in adjusting gene output to cellular needs and are thus critical to animal development and health. Numerous such sequences have been identified in mammalian genomes ranging from transcription factors binding motifs to recognition sites for RNA-binding proteins and non-coding RNAs. The advent of CRISPR has raised the possibility of assigning functionality to individual endogenous regulatory sites by facilitating the generation of isogenic cell lines that differ by a defined set of genetic modifications. Here we investigate the usefulness of this approach to assign function to individual miRNA binding sites. We find that the process of generating isogenic pairs of mammalian cell lines with CRISPR-mediated mutations introduces extensive molecular and phenotypic variability between biological replicates making any attempt of assigning function to the binding site essentially impossible. Our work highlights an important consideration when employing CRISPR editing to characterize non-coding regulatory sequences in cell lines and calls for the development and adoption of alternative strategies to address this question in the future.

## INTRODUCTION

MicroRNAs (miRNAs) are small non-coding RNAs of about 22 nucleotides (nts) in length that direct posttranscriptional repression of mRNA targets by Argonaute proteins [1]. Many miRNAs are highly conserved across species suggesting they play important functions in animal physiology [2]. Indeed, loss-of-function models for many of these genes have uncovered a myriad of phenotypes ranging from embryonic and perinatal lethality to roles in the tumorigenic process [1, 3]. Not surprisingly, miRNA deregulation has often been linked to human disease [4, 5].

Significantly less is known about the functional importance of individual miRNA/mRNA targeting events. MicroRNAs recognize their targets via short sequence complementarity, primarily between the miRNA’s seed (nucleotides 2-8) and the mRNA’s 3’ untranslated region (UTR). The small size of this binding motif means that an individual miRNA can have hundreds of computationally predicted targets, with many being conserved across species [6]. Many of these putative pairing events can also be experimentally captured with strategies such as CLIP [7–10]. Moreover, both computationally and experimentally predicted targets are preferentially upregulated when the cognate miRNA is disrupted [11]. Yet, while these complementary strategies can give insights into the potential targeting repertoire of a miRNA, they do not provide information regarding the functional relevance of individual interactions. In fact, virtually all predicted miRNA targeting events remain functionally unvalidated, with a few notable exceptions [12–18].

Targeted disruption of the miRNA binding site in the endogenous locus of the target gene is currently viewed as the most stringent approach to validate the functional relevance of individual targeting events. Disruption of a functional site is expected to cause not only up-regulation of the target transcript but also the recapitulation—at least partially—of the phenotypes observed in the loss-of-function mutant for the cognate miRNA. Disruption of predicted miRNA binding sites has an additional appeal. Many miRNAs belong to seed-families: groups of miRNAs that share the same seed-sequence and are therefore expected to target largely overlapping sets of transcripts. For many of these miRNAs the extensive targeting redundancy makes it impractical to generate loss-of-function mutants since multiple loci may need to be sequentially mutated before a phenotype becomes apparent. In these cases, disruption of a single binding site that is targeted by all family members may uncover miRNA functions that would be otherwise unknown.

To date, only a few miRNA binding site mutants have been reported [12–18]. The development of CRISPR editing tools, which facilitate the introduction of precise modifications in eukaryotic genomes, raises the possibility of rapidly expanding this number and improving our understanding of both miRNA functions and the common features of functional sites. Particularly, when combined with the ease of cell culture experiments, CRISPR targeting could enable the rapid generation of isogenic cell lines carrying binding site mutations in a variety of cellular pathogenic contexts thus providing a wealth of genetic data as previously done for protein coding genes [19–24]. Here, we investigate the usefulness of this approach using the putatively oncogenic *miR-19*/*Pten* interaction as a model system. We found that isogenic cell lines carrying CRISPR-based *miR-19* binding site mutations displayed molecular and cellular phenotypes that could not be assigned to loss of *Pten* targeting by *miR-19*. Systematic analysis of the source of these signals showed they stemmed not only from the CRISPR targeting procedure itself but also from the clonal expansion required to generate isogenic lines, both of which contributed to extensive variation between clonal replicates. Given the modest effects expected from most miRNA/target interactions, we conclude that this approach is not suitable for their functional dissection. Our work highlights an important consideration when employing CRISPR editing to characterize non-coding regulatory sequences in cell lines and calls for the development and adoption of alternative strategies to address this question in the future.

## MATERIALS AND METHODS

### Cell culture

The MyC-CaP prostate epithelial cell line was purchased from ATCC (CRL-3255) and cultured in 1x DMEM (Gibco, cat# 11965092) supplemented with 10% FBS, L-glutamine (2mM), penicillin (100 U ml^-1^), and streptomycin (100 μg ml^−1^) at 37°C with 5% CO_2_. To generate single cell clones, Myc-CaP cells were trypsinized, pelleted down through centrifugation, and re-suspended in cold PBS containing 2% FBS at approximately 10 million cells ml^-1^. Cells were strained through a nylon mesh (75mm), FACS sorted into individual wells of a 96-well-plate and grown in conditioned media for approximately a week before expanding to a 48-well-plate. Once confluence was reached, half of the cells were used for genotyping and the remaining cells kept in culture for further expansion.

### CRISPR genomic editing

To generate isogenic cell lines carrying targeted genomic mutations, we assembled Cas9/gRNA RNP complexes *in vitro* by mixing 16 μg of recombinant Cas9 (IDT) and 120 pmol of *in vitro* transcribed gRNA (IDT) in 100 μl of SE buffer (Lonza, cat# PBC1-00675) and incubating at room temperature for 10 minutes. For homologous-directed repair, 100 pmol of a DNA oligo corresponding to the template for homologous recombination was added to the mix just before nucleofection. This mixture was used to re-suspend 3 million cells which were electroporated in a Lonza 4D nucleofector using the CM-147 protocol. Cells were plated in conditioned media and sorted into single wells the following day. Sequences of gRNAs used for these experiments are the following: *Pten* site1: CUGUUGCCACAAGUGCAAAG; *Pten* site2: UCAUUGUAAUAGAAUGUGUA and UUAAAUGUCAUUAACUGUUA; *Rosa26* locus: ACTCCAGTCTTTCTAGAAGA. To achieve homologous recombination at Pten site 1 we co-delivered a DNA repair template with the following sequence: ACAGGCTGATGTGTATACGCAGGAGTTTTTCCTTTATTTTCTGTCACCAGCTGAAGTGGCT GAAGAGCTCTGATTCCCGGGTTCACGTCCTACCCCT<u>gaattc</u>TTGTGGCAACAGATAAGTTTG CAGTTGGCTAAGGAAGTTTCTGCAGGGTTTTGTTAGATTCTAATGCATGCACTTGGGTTGGGAATGGAGGGAATGC, where the small caps underlined sequence corresponds to the replacement of the miR-19 binding site with an EcoRI restriction motif.

For detection of genomic editing, cells were collected in lysis buffer (10% SDS, 5 mM EDTA, 200 mM NaCl, 100 mM Tris pH 8.0) together with 0.2 mg/mL Proteinase K and incubated at 55°C for 1 hour and 800 rpm. Genomic DNA was extracted with phenol-chloroform followed by ethanol precipitation and amplified by PCR using Phusion^TM^ High-Fidelity DNA Polymerase enzyme (Thermo Fisher Scientific, cat# F530). Detection of genomic integration at *miR-19* binding site 1 was achieved by amplifying genomic DNA using primers outside the homology arms to the repair template (forward: CCACAGGGTTTTGACACTTG; reverse: CCAGAGCCCAGGTAGAAACA) resulting in an amplicon of 385 bp. Digestion of this fragment with EcoRI produced fragments of 254 bp and 131 bp in clones that underwent successful recombination. Detection of genomic deletion at *miR-19* binding site 2 was achieved using primers flanking the two gRNA target sites (forward: CTGTGGATGCTTCATGTGCT; reverse: TGAAGCCCTAATCCCAACTC). This resulted in an amplicon of 482 bp in the parental cell line and approximately 420 bp in clones carrying a deletion. Detection of genomic editing at the *Rosa26* locus was performed using primers flanking the gRNA target site (forward: GCTCAGTTGGGCTGTTTTGGAG; reverse: CCAGATGACTACCTATCCTCCCA), which amplifies a fragment of 428 bp, followed by generation of heteroduplexes and digestion of the DNA with the mismatch-sensitive T7 endonuclease.

### RNA isolation and real-time quantitative PCR

Total RNA was isolated from cells using TRIzol (Invitrogen, cat# 15596018) following the manufacturer’s protocol. Isolated RNA was treated with ezDNase (Thermo Fisher Scientific, cat# 18091150) to remove contamination of genomic DNA and reverse transcribed using SuperScript^TM^ IV Reverse Transcriptase (Thermo Fisher Scientific, cat# 18090010) using Oligo-dT primers. Relative expression levels of *Pten* and *Hprt1* were quantified using TaqMan Gene Expression Assays (Thermo Fisher Scientific, *Pten*, Mm00477208_m1; *Hprt1*, Mm00446968_m1) together with TaqMan Fast Advanced Master Mix (Thermo Fisher Scientific cat# 4444557).

### RNA-sequencing data and analysis

RNA sequencing data from *miR-19*-null animals was previously published [25] and can be retrieved from the GEO database (GSE63660). Processed read counts from different tissues were used to identify differentially expressed genes between wild-type and *miR-19*-null animals using DESeq2 [26]. Predicted target genes carrying *miR-19* binding sites were obtained from TargetScanMouse Release 8.0 [11, 27]. Data from MYC-driven tumors was generated in [28].

### Protein isolation and western blot

Cells were washed in PBS and resuspended in 200 μL of RIPA lysis buffer (0.1 % SDS, 50 mM Tris pH 8.0, 1 % NP-40, 0.5 % Sodium deoxycholate, 150 mM NaCl) along with protease inhibitor cocktail (Roche). Protein lysates were quantified using Pierce™ Rapid Gold BCA Protein Assay Kit (Thermo Fisher Scientific, cat# A53225). 25 μg samples were run on 4–12% Bis-Tris precast gels (Invitrogen) and transferred to a nitrocellulose membrane. Membranes were blocked in 5 % milk for 1 hour followed by overnight incubation with PTEN primary antibody (Cell Signaling Technology, cat# 9188; 1:1000 dilution) or GAPDH primary antibody (Santa Cruiz Biotechnology, cat# sc32233; 1:1000 dilution) at 4°C on a rocker. After multiple washes with 1x PBST, the membranes were incubated with respective secondary antibody at 1:5000 dilution for 1 hour at room temperature on a rocker. Membranes were washed three times with 1x PBST and detection was performed on Amersham hyperfilms using ECL^TM^ Detection Reagent (Cytiva, cat# RPN2105).

### Caspase activity quantification

Equal number of cells from each line was seeded on 6-well plates and grown overnight. Next day, cells were trypsinized, pelleted down through centrifugation, and re-suspended in 300 μL of culture media containing 1 μL of FITC-VAD-FMK (Calbiochem Caspase Detection Kit, cat# QIA90) and incubated for 45 minutes in a 37°C incubator with 5 % CO_2_, following manufacturer’s instructions. Cells were pelleted down through centrifugation and washed twice with Wash Buffer. Finally, the cells were re-suspended in 500 μL of Wash Buffer and strained into 5 mL Falcon Polystyrene Round-Bottom Tube with Cell-Strainer Cap (75 mm) for flow cytometric analysis on FACSCanto II (BD Biosciences). FACS data were analyzed with FlowJo^TM^ v10.10.

## RESULTS AND DISCUSSION

### Generation of miR-19 binding site mutants in *Pten*’s 3’UTR

The *miR-17∼92* polycistronic miRNA cluster is a transcriptional target of MYC [29] and is frequently amplified in human tumors [30]. It encodes 6 distinct miRNAs [31]. Two of these (*miR-19a*, *miR-19b-1*) belong to *the miR-19* seed-family and act as key mediators of MYC’s oncogenic properties *in vivo* [25]. Deletion of *miR-19a* and *miR-19b-1* in *Eμ-Myc* mice (a model of human B-cell lymphomas) or in *Hi-Myc* mice (a model of prostate adenocarcinomas) is enough to compromise the tumorigenic process. In both cases, tumor initiation and progression are impaired due to a dramatic increase in cellular apoptosis in response to oncogenic levels of MYC. These pro-survival functions of *miR-19* can also be observed *in vitro*. In established MYC-driven tumor cell lines, deletion of *miR-17∼92* causes a significant increase in cell death, which can be rescued by re-introduction of *miR-19* alone [28]. This phenotype causes cells lacking *miR-19* to be out-competed by those with an intact locus over the course of a few days. Thus, this *in vitro* system represents a perfect setting to test our ability to assign function to predicted miRNA/mRNA interactions.

While the molecular mechanisms underlying the pro-survival functions of *miR-19* are not well understood, *Pten* has been extensively implicated as one of its functional targets. *Pten* contains two binding sites for *miR-19* in its 3’UTR, a 7mer-m8 (site 1, s1) and an 8mer (site 2, s2) (**Figure 1A**). Both these binding sites are highly conserved across species (**Supplementary Figure 1A**) suggesting they are physiologically important. In line with this, loss of *miR-19* results in up-regulation of *Pten* levels and silencing *Pten* with shRNAs in *miR-17∼92*-null tumors can partially suppress the apoptotic phenotype in these cells [28]. *Pten* is a well-established tumor suppressor gene displaying haploinsufficient phenotypes upon heterozygous loss [32, 33], indicating that even minor changes in its expression—as those observed upon loss of miRNA targeting—can have physiological consequences. Given that increased PTEN levels can trigger apoptosis [34], the emerging model is that in MYC-driven tumors, transcriptional up-regulation of *miR-1*9 promotes cell survival by curtailing the levels of *Pten* and downstream apoptosis, therefore enabling tumor growth (**Figure 1B**).

**Figure 1.**
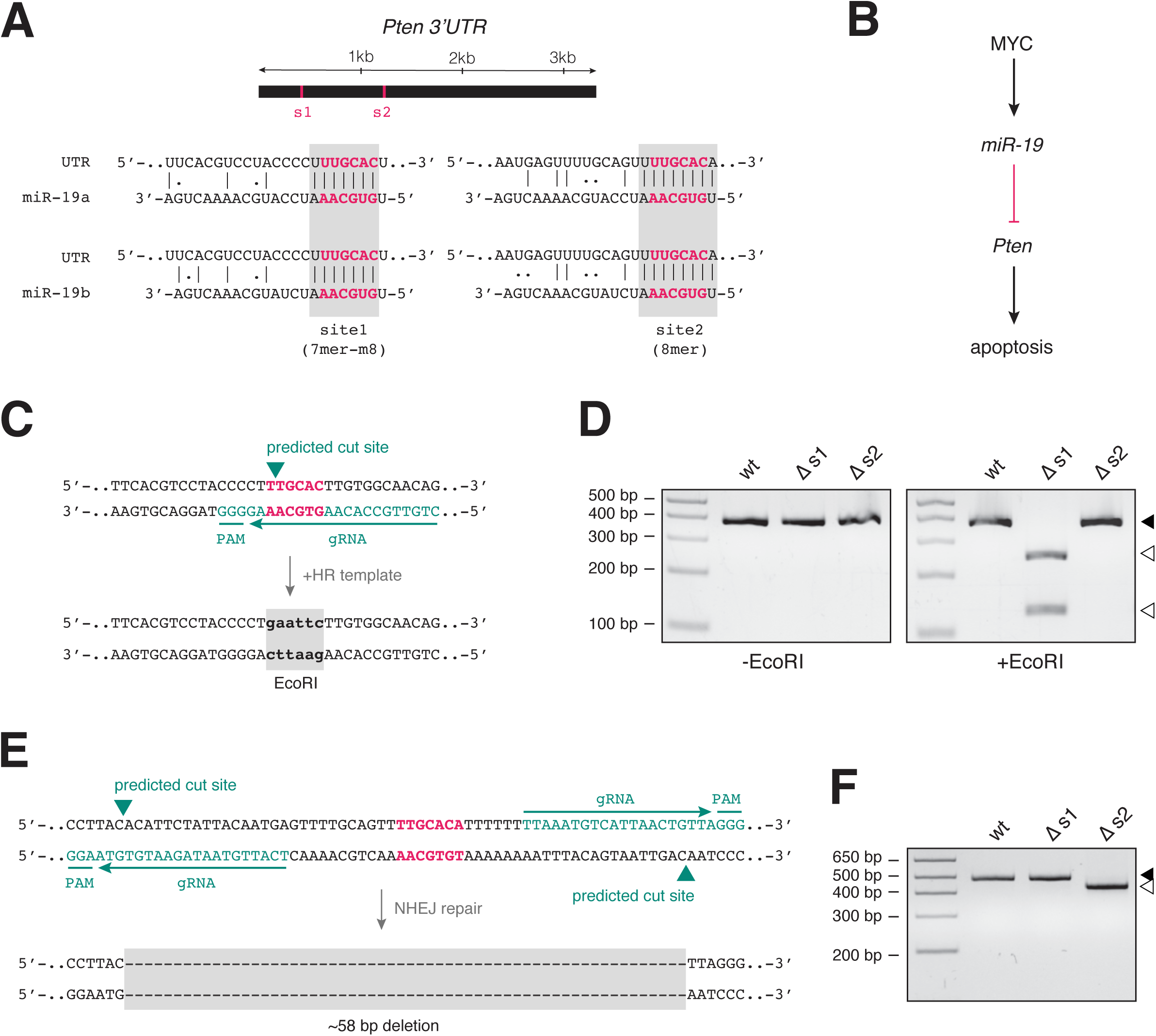
Generation of isogenic cell lines carrying targeted disruptions of *miR-19* binding sites in *Pten*’s 3’UTR. **(A)** A schematic representation of *Pten*’s 3’UTR is shown on top, with the position of the two *miR-19* binding sites (s1 and s2) highlighted. The sequence of each binding site as well as its complementarity to *miR-19a* and *miR-19b* is shown below. Dashes represent standard Watson-Crick matches. Dots represent G:U wobble pairings. The seed sequence on the miRNAs and the seed-match on the 3’UTR are highlighted in bold and color. **(B)** Schematic representation of the MYC-*miR-19*-*Pten* axis and its impact in cell survival. **(C)** Targeting strategy to disrupt binding site 1 (*Δs1*) through homologous recombination (HR). Correct repair results in the replacement of the seed-match sequence with an EcoRI restriction site. Position of the gRNA, PAM sequence, and predicted cut site are highlighted. **(D)** Example of genotyping gel for binding site 1 mutation (*Δs1*), showing EcoRI digestion products in a binding site 1 mutant line (*Δs1*) but not in the parental line (wt) or in a line carrying a targeted disruption of binding site 2 *(Δs2*). Closed arrowhead shows the position of undigested amplicons. Open arrowheads show the position of the digestion products. **(E)** Targeting strategy to disrupt binding site 2 through a targeted deletion. Non-homologous end joining (NHEJ) mediated repair of the DNA breaks following two simultaneous Cas9-mediated cuts results in a deletion of approximately 58 bp encompassing the binding site. **(F)** Example of genotyping gel for binding site 2 mutation, showing the deleted allele only in a line carrying a targeted disruption of binding site 2 (*Δs2*). Closed arrowhead shows the position of the wild-type amplicon. Open arrowhead shows the position of the amplicon carrying the targeted deletion.

To provide genetic support for this model we set out to disrupt each of these binding sites *in vitro* and test the effect of those modifications on both *Pten* levels and cell survival. We used MyC-CaP cells as a model system, as these are derived from the Hi-Myc prostate model where we have previously shown *miR-19* plays an oncogenic role [25]. To minimize variation in *Pten* expression in subsequent experiments due to initial cell line heterogeneity we first established a parental line clonally expanded from a single cell. Next, to mutate binding site 1 we designed a gRNA predicted to cut within the seed-match sequence and co-delivered a repair template that would replace that binding site with an EcoRI recognition motif (**Figure 1C**). Amplification of the locus with primers outside the homology region, followed by EcoRI digestion showed correct homozygous integration of the repair template by homologous recombination and generation of binding site 1 mutant alleles *(Δs1*; **Figure 1D**). This was further confirmed by sequencing of the amplicons (**Supplementary Figure 1B**). Because of the lack of suitable gRNAs, we were unable to follow the same strategy for binding site 2. Instead, we designed and delivered two gRNAs flanking the seed-match sequence. Repair of the double-stranded breaks by non-homologous end-joining (NHEJ) resulted in homozygous deletion of about 58 base pairs (bp) around the binding site *(Δs2*) (**Figure 1E**), which we confirmed by both PCR (**Figure 1F**) and sequencing (**Supplementary Figure 1C**).

### Isogenic cell lines carrying *miR-19* binding site mutations display extensive phenotypic variation

Transcript targeting by miRNAs results in the recruitment of a large riboprotein complex (RISC) that ultimately leads to repression of gene expression primarily through mRNA destabilization. As a consequence, loss of a miRNA or disruption of its target site is expected to result in a significant but mild up-regulation of the targeted mRNA. Accordingly, RNA-sequencing of animals carrying a targeted deletion of *miR-19* showed a significant but modest up-regulation of predicted target transcripts—defined here as those carrying conserved 8-mer binding sites for *miR-19* in their 3’UTRs—compared to wild-type controls, even with increasing numbers of binding sites (**Figure 2A**). Moreover, the expression of *Pten* itself, one of the best characterized targets of *miR-19*, showed only mild up-regulation in the absence of *miR-19* (**Figure 2B**), both in embryonic tissues (mean upregulation of 1.2-fold) and MYC-driven tumors (mean upregulation of 1.3-fold).

**Figure 2.**
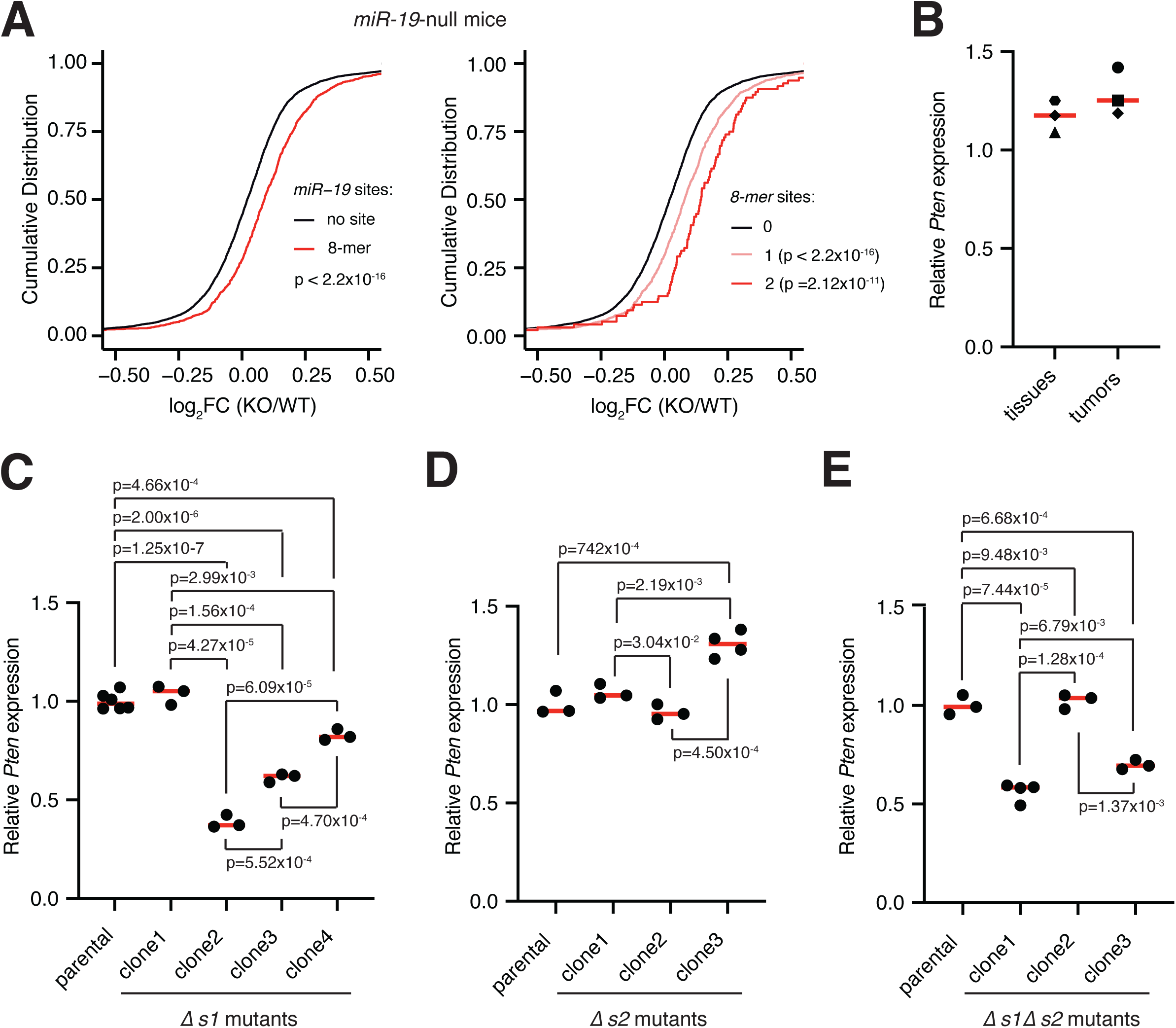
Characterization of *Pten* levels in *miR-19* binding site mutants. **(A)** Cumulative distribution function (CDF) plot showing deregulation of gene expression in animals lacking *miR-19* versus wild-type controls. Left, CDF plot for mRNAs with no predicted binding site for *miR-19* (black) versus those containing one or more 8-mer binding sites (red). Right, CDF plot for mRNAs with no predicted binding site for *miR-19* (0, black), versus those with one (1, light red) or two (2, dark red) 8-mer sites. P values were calculated with the Kolmogorov–Smirnov test. **(B)** Mild upregulation of *Pten* expression upon genetic deletion of *miR-19*. Each data point represents the mean of three biological replicate samples collected from different embryonic tissues or tumor cells. Red bar shows mean value of tissue or tumor data. **(C)** Relative *Pten* expression in clonal cell lines carrying a homozygous disruption of binding site 1 *(Δs1*) compared to the levels in the parental cell line. **(D)** As in (A) but for binding site 2 mutants (*Δs2*). (E) As in (A) but for cells carrying homozygous disruption of both binding sites *(Δs1Δs2*). For all plots, p values were calculated with a two-tailed t-test. Each dot represents a technical replicate. Red bars indicate mean of replicates.

To determine how disruption of *miR-19* binding motifs on *Pten*’s transcript compared to these changes after loss of *miR-19*, we measured *Pten* expression in our binding site mutant cell lines relative to the parental control. Surprisingly, we found that isogenic clonal cell lines carrying targeted disruptions of binding site 1 (*Δs1*; **Figure 2C**) or binding site 2 (*Δs2*; **Figure 2D**) showed varying *Pten* levels relative to the control line, with extensive and significant variation between clonal replicates. This variation was also present in double mutant cell lines *(Δs1Δs2*) carrying concomitant disruption of both motifs (*Δs1Δs2*; **Figure 2E**). Importantly, the magnitude of variation observed among all clonal lines was within, and frequently beyond, the ranges expected upon loss of miRNA targeting, as the magnitude of changes observed in *miR-19* loss-of-function samples relative to controls were substantially smaller than those we observed in binding site mutant lines (**Figure 2B-E**). Moreover, contrary to the predicted effect of miRNA binding site disruption, we found that *Pten* expression was surprisingly often lower in mutant lines compared to the parental control.

Transcript targeting by miRNAs ultimately results in down-regulation of protein abundance. Although a component of translational inhibition has been documented, changes in protein levels are for the most part secondary to the destabilization of the target mRNA and therefore closely reflect transcript levels [35]. To determine how PTEN levels changed upon *miR-19* binding site mutation, we performed western blot analysis to all clonal lines compared to the parental cell line. As before, we observed extensive variation between biological replicates (**Figure 3A**). Importantly, we found poor correlation between mRNA and protein levels or the number of mutated binding sites (**Figure 3B**). This suggests that the increase in protein abundance is independent of *miR-19*’s effect on mRNA stability.

**Figure 3.**
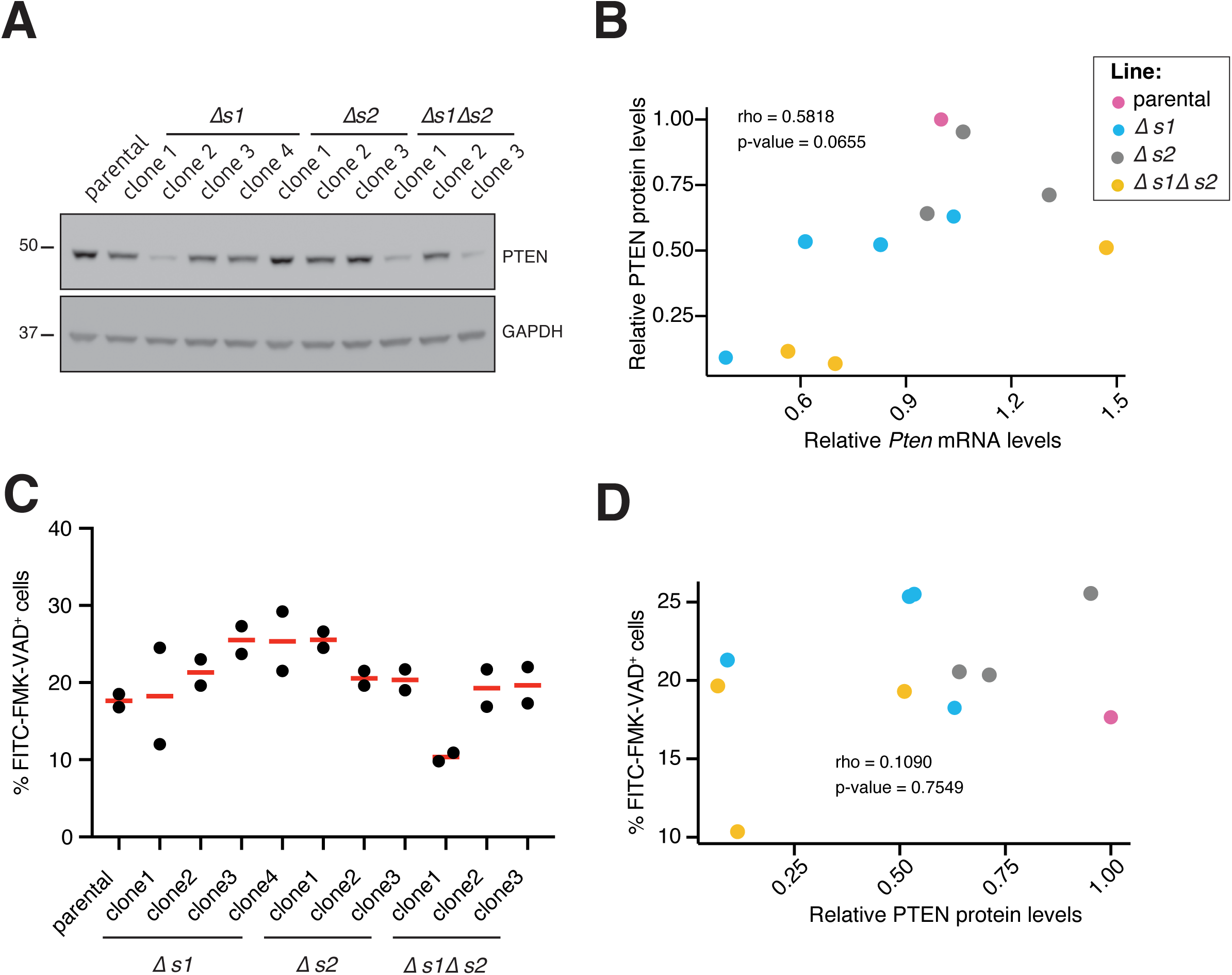
Characterization of PTEN and activated caspase levels in *miR-19* binding site mutants. **(A)** Western blot to PTEN in binding site mutants and control parental cell line. **(B)** Scatter plot comparing relative levels of *Pten* mRNA and protein in binding site mutant lines compared to parental control. The Pearson correlation coefficient (rho) between the two variables, as well as the significance of that correlation are shown. **(C)** Caspase activity in binding site mutant cell lines as measured by flow cytometry using FITC-VAD-FMK. The percentage of FMK positive cells for each cell line is shown. **(D)** As in (B) but comparing relative PTEN protein levels with caspase activity levels for all lines.

Finally, we examined the phenotypic consequences of mutating *miR-19*’s binding sites on *Pten*’s 3’UTR. Specifically, we measured the fraction of cells undergoing apoptosis by detecting caspase activation through flow cytometry (**Figure 3C**, **Supplementary Figure 2**). Since repression of *Pten* by *miR-19* is thought to be required for survival of MYC-driven tumor cells, disruption of this regulatory interaction is predicted to result not only in higher expression of this tumor suppressor gene, but also in higher levels of apoptosis. While we did observe a modest increase in activated caspase in many mutant clones, these changes were again highly variable between replicate lines (**Figure 3C**) and importantly, showed no correlation with PTEN protein levels (**Figure 3D**), suggesting the two are unrelated.

Taken together, our molecular and phenotypic analysis of multiple clonal lines carrying CRISPR-derived *miR-19* binding site mutations suggests that the majority of changes observed are not a direct consequence of disrupting the miRNA/target interaction. Instead, we speculated that they might arise from the targeting protocol itself, which we investigate further below.

### Clonal variability following CRISPR targeting and clonal expansion

To determine the source of variation observed among binding site mutant clones, we re-targeted the MyC-CaP parental cell line with a gRNA designed to disrupt the *Rosa26* locus with high specificity (**Figure 4A, 4B**) [36]. We selected this locus because *Rosa26* has been frequently used as a location for the expression of transgenes in the mouse because its disruption does not have a significant impact on animal development or health [37]. As a consequence, disruption of Rosa26 with CRISPR should have no significant impact on *Pten* expression or cell survival. We expanded 10 independent clones following targeting which we used to characterize how disruption of a safe-harbor locus with CRISPR affected *Pten* levels and cell viability. In parallel, we expanded 10 additional clones that were not subject to targeting to determine the consequences of the clonal expansion process itself. Surprisingly, we found that *Rosa26* targeting generated extensive variation in *Pten* mRNA levels amongst clones, more than half of which showed significant differences to the parental line (**Figure 4C**). Remarkably, even the simple process of clonal expansion itself was sufficient to generate these high levels of clonal variation (**Figure 4D**).

**Figure 4.**
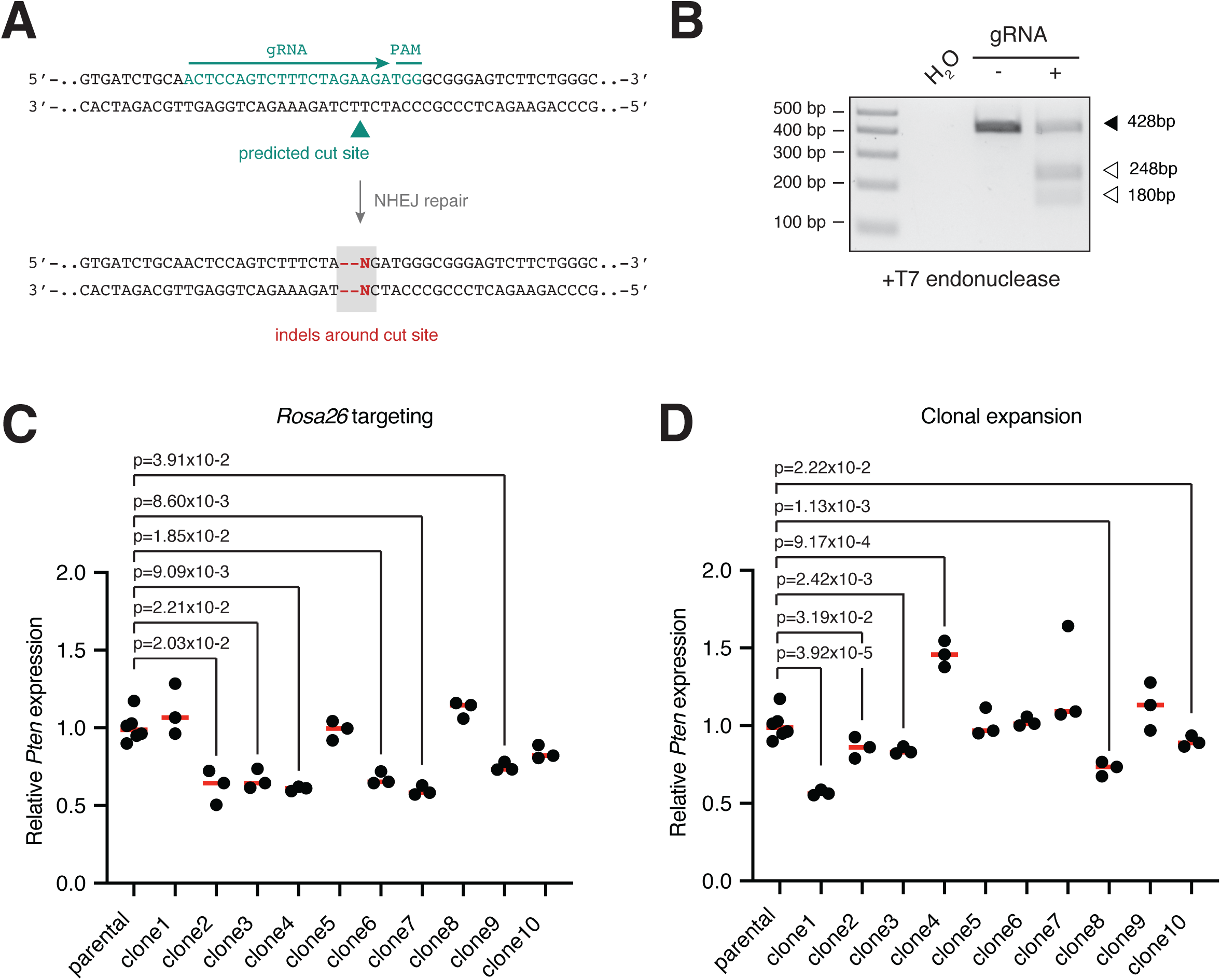
Clonal variation in *Pten* levels following CRISPR targeting and clonal expansion. **(A)** Targeting strategy for the *Rosa26* locus. Position of the gRNA, PAM sequence, and predicted cut site are highlighted. Repair of the DNA break by Non-homologous end joining (NHEJ) results in the generation of indels around the cut site. **(B)** Detection of indels in cells targeted with a gRNA against the *Rosa26* locus using the mismatch-sensitive T7 endonuclease. Digestion of heteroduplexes by T7 leads to the generation of two cleavage products (open arrowheads) of the indicated sizes. Position of undigested amplicons is shown by the closed arrowheads. **(C)** Relative expression levels of *Pten* in clonal cell lines expanded following the targeting of the *Rosa26* locus compared to the levels in the parental cell line. P value, two-tailed t-test. Each dot represents a technical replicate. Red bars indicate mean of replicates. **(D)** As in (C) but for single cell clones not subjected to CRISPR targeting.

To extend these observations to other phenotypes linked to the *miR-19*/*Pten* axis, we next checked how targeting of the safe-harbor locus, or the process of clonal expansion affected PTEN protein levels or cell viability. Similarly to what we observed with binding site mutant lines, both clones derived from the *Rosa26* targeting (**Figure 5A**) and those obtained following simple clonal expansion (**Figure 5B**) tended to show significantly higher protein levels than the parental line, with no correlation with the measured transcript abundance (**Figure 5C**, **Figure 5D**). As before, we also measured a significant variability in caspase activity levels in clonal lines from both experiments (**Figure 5E**, **Figure 5F**, **Supplementary Figure 3A**, **Supplementary Figure 3B**), and found no correlation between these levels and those of PTEN (**Figure 5G**, **Figure 5H**). These data indicate that the simple process of generating isogenic lines following CRISPR targeting is enough to create substantial variation between clones across multiple aspects of cell biology.

**Figure 5.**
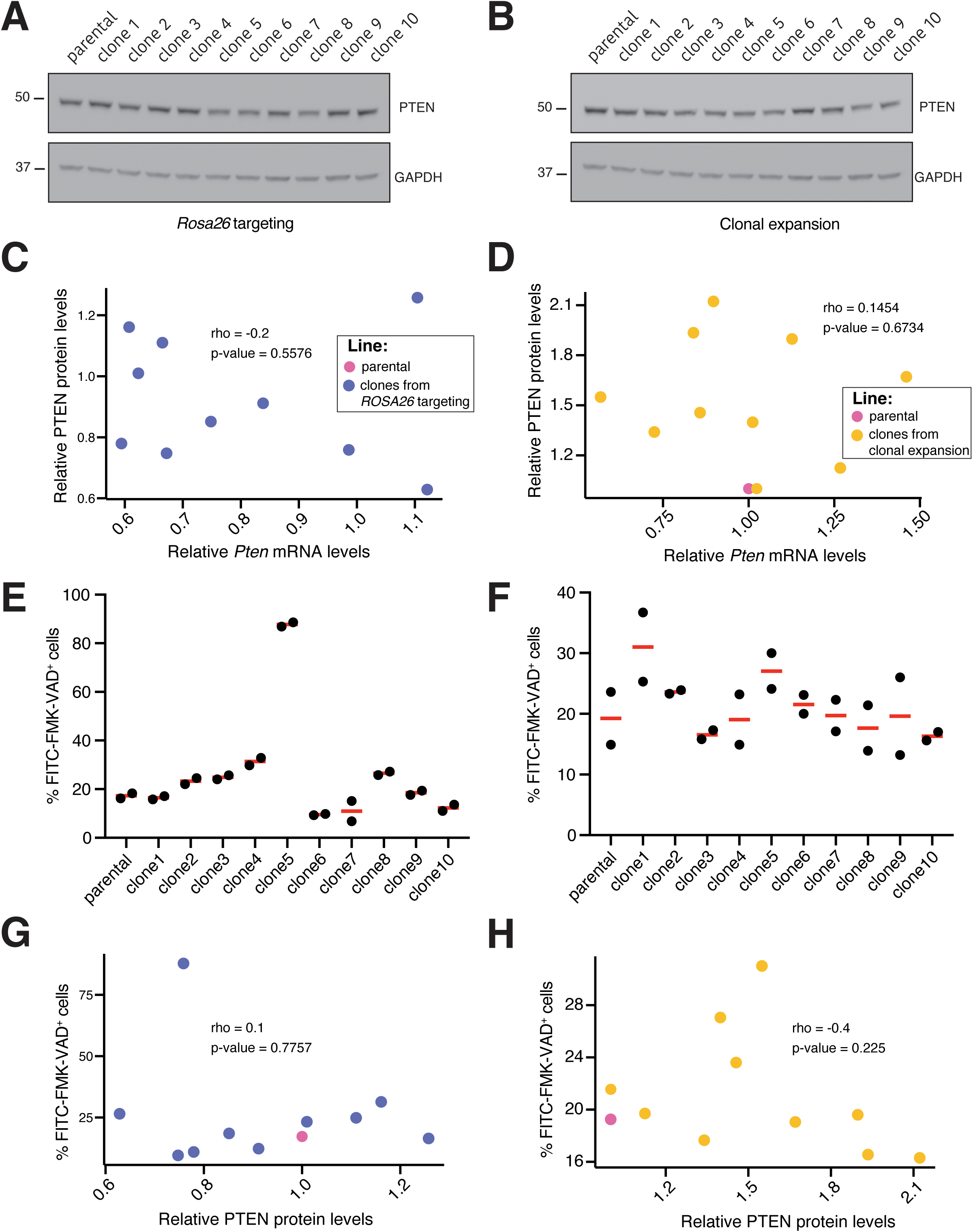
Variable PTEN levels and apoptosis in isogenic lines following CRISPR targeting and clonal expansion. **(A)** Western blot to PTEN in clones derived from *Rosa26* targeting and control parental cell line. **(B)** Western blot to PTEN in clones derived from simple clonal expansion compared to parental control. **(C)** Scatter plot comparing relative levels of *Pten* mRNA and protein in clones derived from *Rosa26* targeting (purple) compared to parental control (pink). The Pearson correlation coefficient (rho) between the two variables, as well as the significance of that correlation are shown. **(D)** As in (C) but for lines derived from clonal expansion (green) compared to parental control (pink). **(E)** Caspase activity in *Rosa26* targeting clones as measured by flow cytometry using FITC-VAD-FMK. The percentage of FMK positive cells for each cell line is shown. **(F)** As (E) but for lines derived from clonal expansion. **(G)** As in (C) but comparing relative PTEN protein levels with caspase activity levels for *Rosa26* targeting lines lines. **(H)** As in (G) but for lines derived from clonal expansion.

Isogenic lines that differ from one another by targeted genomic modifications have been instrumental tools to assign gene functions, particularly for genomic alterations that impact cancer development. Initially, such lines were generated using traditional gene targeting techniques which are both time-consuming and technically challenging to implement, essentially limiting this approach to a limited number of researchers, areas of study, and to protein coding genes. The development of CRISPR as a genome editing tool has overcome many of the technical challenges underlying the generation of isogenic lines making their implementation accessible to a larger number of researchers. Although generation of isogenic cells remains time consuming, it can now be conceivably applied to query the functional relevance of essentially any genomic sequence including non-coding regulatory regions such as UTRs [38] or enhancer sequences [39].

The use of isogenic lines to study the function of miRNA binding sites is particularly appealing. Validation of these sites on potential target transcripts is typically done using reporter assays in which wild-type or mutated motifs are inserted directly downstream of a transgene whose expression can be easily measured following transfection. Yet, given the known impact of the miRNA/target stoichiometry for effective repression, these assays can easily fail to identify true targets. The generation of isogenic cells carrying disrupted binding site motifs circumvents these issues by testing the levels of de-repression of the putative target under the physiological levels of both its transcript and the targeting miRNA. Moreover, this data can be theoretically complemented with phenotypical assays to determine the extent to which binding site disruption re-capitulates the loss of the cognate miRNA. Although still relatively time consuming, this approach promised to provide a more robust understanding of miRNA biology [1].

Following the design and delivery of the targeting reagents, the generation of engineered isogenic lines requires the isolation and expansion of single cells, which can then be characterized for the correct genotype and downstream molecular and cellular phenotypes. While employing this approach to genetically validate the well-studied *miR-19*/*Pten* pro-survival interaction in MYC-driven tumors, we found that the process of clonal expansion alone resulted in levels of clonal variation extensive enough to essentially preclude us from determining the functional impact of *miR-19* binding site disruption on target expression and cell fitness. We conclude, that genetically engineered isogenic cells, which have proven so useful for the study of protein coding genes [19–24], are not compatible with the study of miRNA biology. Indeed, the modest repression effect expected from most interactions is likely to be masked by the cellular changes acquired during the clonal expansion process.

Given these issues, the genetic validation of miRNA/target interactions will likely need to rely on alternative strategies that circumvent the need to generate single cell clones. One such strategy is the generation of genetically engineered animals carrying the desired mutation. This has already been successfully applied to the validation of a few target sites [12–18] and has the additional strength of yielding physiologically relevant data over multiple cellular contexts. Yet, the significant investment this approach requires combined with its low throughput means it will likely be adopted by only a handful of labs and used to validate only the most promising interaction pairs. As a result, potential insights into novel miRNA biology that could result from studying less canonical sites (for example those that are not conserved across species or that do not contain a seed-match) is likely to be missed. Alternatively, functional validation of binding sites could be performed using high-throughput CRISPR screens, which enable large-scale testing of loci without the need to expand clonal lines. The employment of screens to study individual miRNA/target interactions in a high-throughput manner will likely require the development of tools to deal with the low available targeting space, exemplified here by our inability to target binding site 2 with a single specific gRNA. Given that comprehensive targeting of binding sites is incompatible with stringent selection of highly-specific gRNAs, tools that can correct for the effects of off-targeting [40] may be particularly useful to these efforts.

In summary, our work highlights the need to develop specific strategies for the study of miRNA regulatory sites and likely other non-coding regulatory regions expected to impart subtle changes on gene expression. By identifying key obstacles posed by current protocols, we hope this study will help in the development of such strategies, more suitable to the study of these loci.

## Supporting information

SupFigure1

SupFigure2

SupFigure3

## DATA AVAILABILITY

RNA sequencing data from *miR-19*-null animals was previously published [25] and can be retrieved from the GEO database (GSE63660)

## FUNDING

This work was supported by the Intramural Research Program of the National Institutes of Health through the Center for Cancer Research, National Cancer Institute, project 1ZIABC011810-02 (JAV) and a FLEX grant from the Center for Cancer Research (JAV).

## ACKNOWLEDGEMENTS

We thank members of the Vidigal lab for discussions and comments on this work. We thank the CCR/LGI Flow Cytometry Core, particularly Ferenc Livák, Shafiuddin Siddiqui, and Karen M. Wolcott for their outstanding technical support. This work utilized the computational resources of the NIH HPC Biowulf cluster (hpc.nih.gov).

## SUPPLEMENTARY FIGURE LEGENDS

**Supplementary Figure 1. *miR-19* binding sites in *Pten*’s 3’UTR. (A)** Sequences on *Pten*’s 3’UTR with complementarity to *miR-19* across species (left, binding site 1; right, binding site 2). Seed match regions are highlighted with bold characters and grey box. Nucleotides in which the sequence in other species differs from that of mice are highlighted in color. **(B)** Example sanger sequencing traces showing correct integration of HDR donor template in binding site 1 locus. The position of the EcoRI site is highlighted. **(C)** Example sanger sequencing traces showing deletion of binding site 2. The position of the PAMs for both targeting gRNAs as well as the position of the repair following the DNA breaks are shown.

**Supplementary Figure 2. Generation of isogenic cell lines carrying targeted disruptions of *miR-19* binding sites in *Pten*’s 3’UTR. (A)** Example flow cytometry histograms of caspase activity as measured by percentage of VAD-FMK^+^ cells in the parental control line and in biological triplicate lines carrying single *(Δs1*, *Δs2*) or double (Δs1Δs2) binding site mutations.

**Supplementary Figure 3. Generation of isogenic cell lines carrying targeted disruptions of *miR-19* binding sites in *Pten*’s 3’UTR. (A)** Example flow cytometry histograms of caspase activity as measured by percentage of VAD-FMK^+^ cells in the parental control line and in the 10 clones derived from the *Rosa26* targeting. **(B)** As in (A) but for 10 cell lines derived from simple clonal expansion.

## Notes

### Competing Interest Statement

The authors have declared no competing interest.

