## Supplementary material for "CRISPR-based dissection of miRNA binding sites using isogenic cell lines is hampered by pervasive noise": SupFigure1

### Prajapat, Sup. Figure 1

**A**

| | site1 | site2 |
|---|---|---|
| mouse | 5′–..UUCACGUCCUACCCCU**UUGCACU**..–3′ | 5′–..AAUGAGUUUUGCAGUU**UUGCACA**..–3′ |
| rat | 5′–..UUCACGUCCUACCCCU**UUGCAUU**..–3′ | 5′–..AAUGAGUUUUGCAGUU**UUGCACA**..–3′ |
| guinea pig | 5′–..UUCACAUCCUACCCCU**UUGCACU**..–3′ | 5′–..AAUGAGUUUUGCAGU–**UUGCACA**..–3′ |
| squirrel | 5′–..UUCACAUCCUACCCCU**UUGCACU**..–3′ | 5′–..AAUGAGUUUUGCAGUU**UUGCACA**..–3′ |
| rabbit | 5′–..UUCACAUCCUACCCCU**UUGCACU**..–3′ | 5′–..AAUGAGUUUUGCAGUU**UUGCACA**..–3′ |
| pika | 5′–..UUCAUAUCCUACCCCU**UUGCACU**..–3′ | 5′–..AAUGAGUUUUGCAGUU**UUGCACA**..–3′ |
| human | 5′–..UUCACAUCCUACCCCU**UUGCACU**..–3′ | 5′–..AAUGAAUUUUGCAGUU**UUGCACA**..–3′ |
| rhesus | 5′–..UUCACAUCCUACCCCU**UUGCACU**..–3′ | 5′–..AAUGAGUUUUGCAGUU**UUGCACA**..–3′ |
| chinese pangolin | 5′–..UUCACAUCCUACCCCU**UUGCACU**..–3′ | 5′–..AAUGAGUUUUGCAGUU**UUGCACA**..–3′ |
| dog | 5′–..UUCACAUCCUACCCCU**UUGCACU**..–3′ | 5′–..AAUGAGUUUUGCAGUU**UUGCACA**..–3′ |
| sheep | 5′–..UUCACAUCCUACCCCU**UUGCACU**..–3′ | 5′–..AAUGAGUUUUGCAGUU**UUGCACA**..–3′ |
| opossum | 5′–..CUCCCGGCCUGCCCC–**UCGCACU**..–3′ | 5′–..CAUGAGUUUUGCAGUU**UUGCACA**..–3′ |
| | seed match | seed match |

**B**

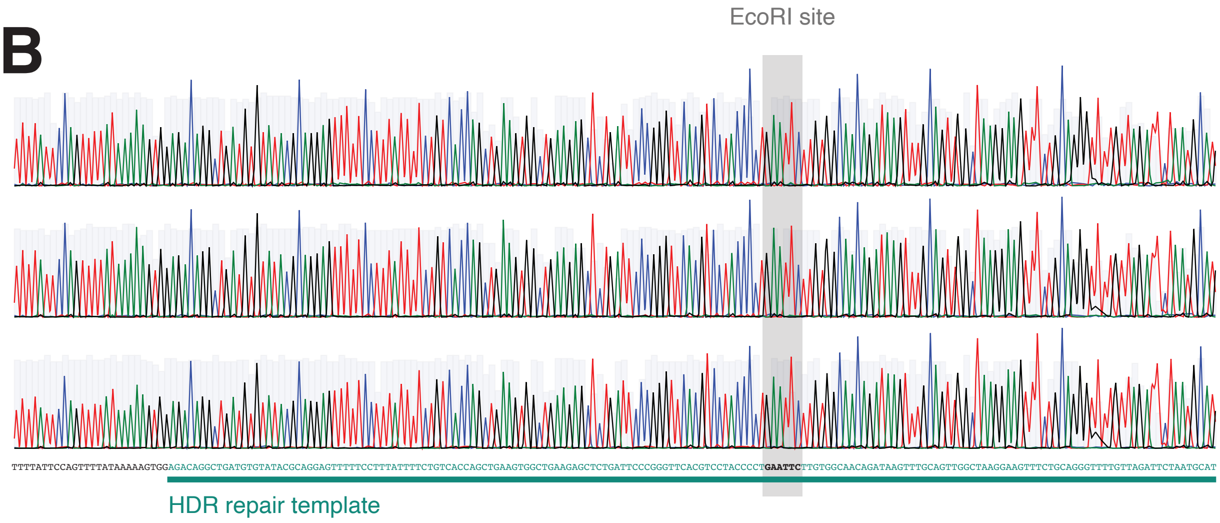


**C**

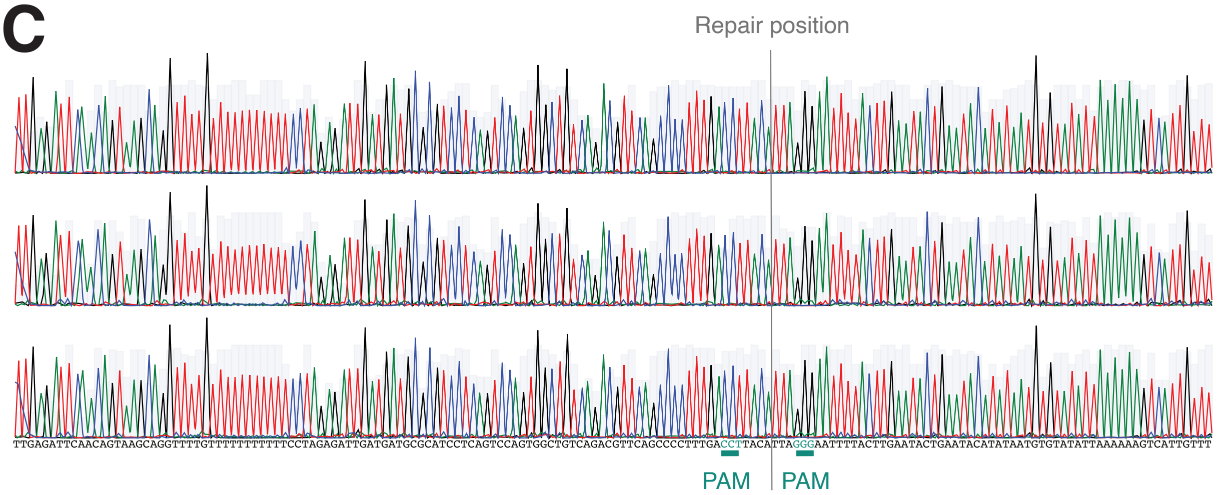
