## Supplementary figures and images for "CRISPR-based dissection of miRNA binding sites using isogenic cell lines is hampered by pervasive noise"

### SupFigure2

#### Prajapat, Sup. Figure 2

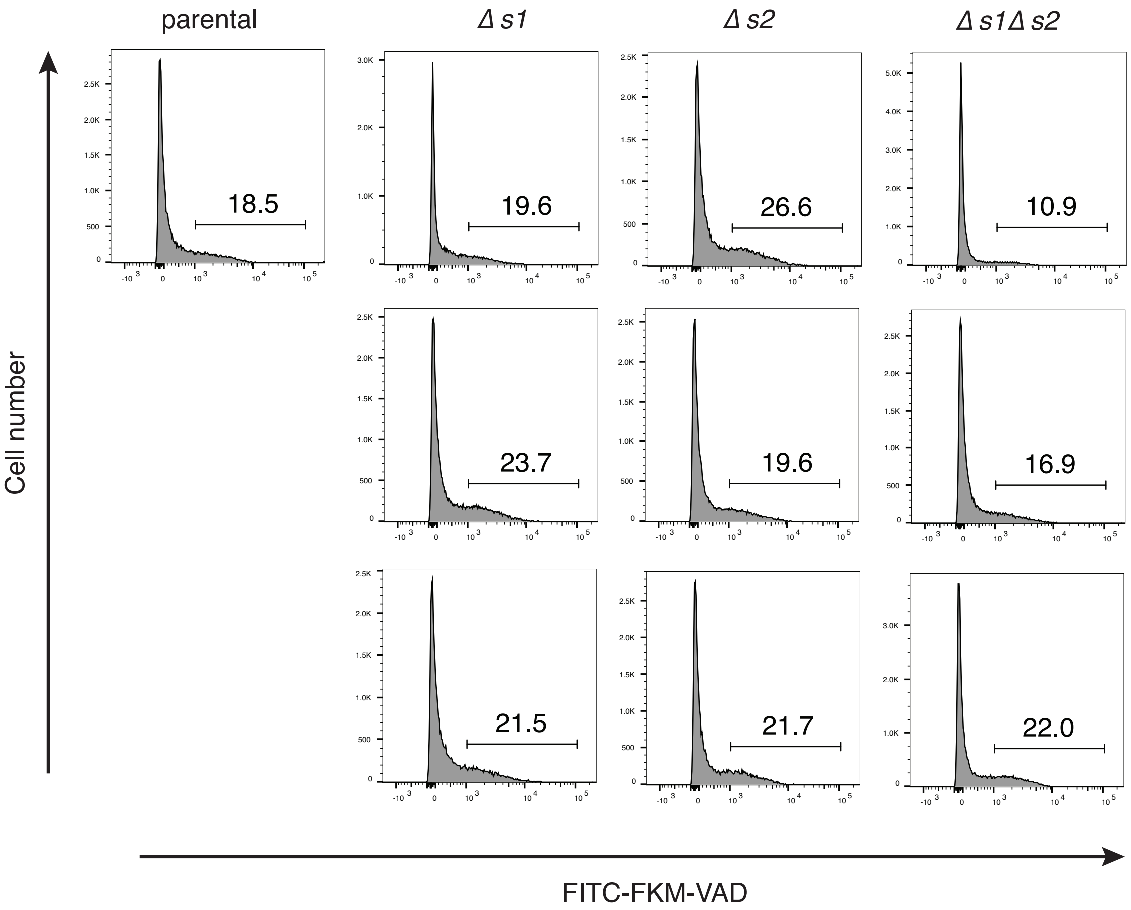

### SupFigure3

**Prajapat, Sup. Figure 3**

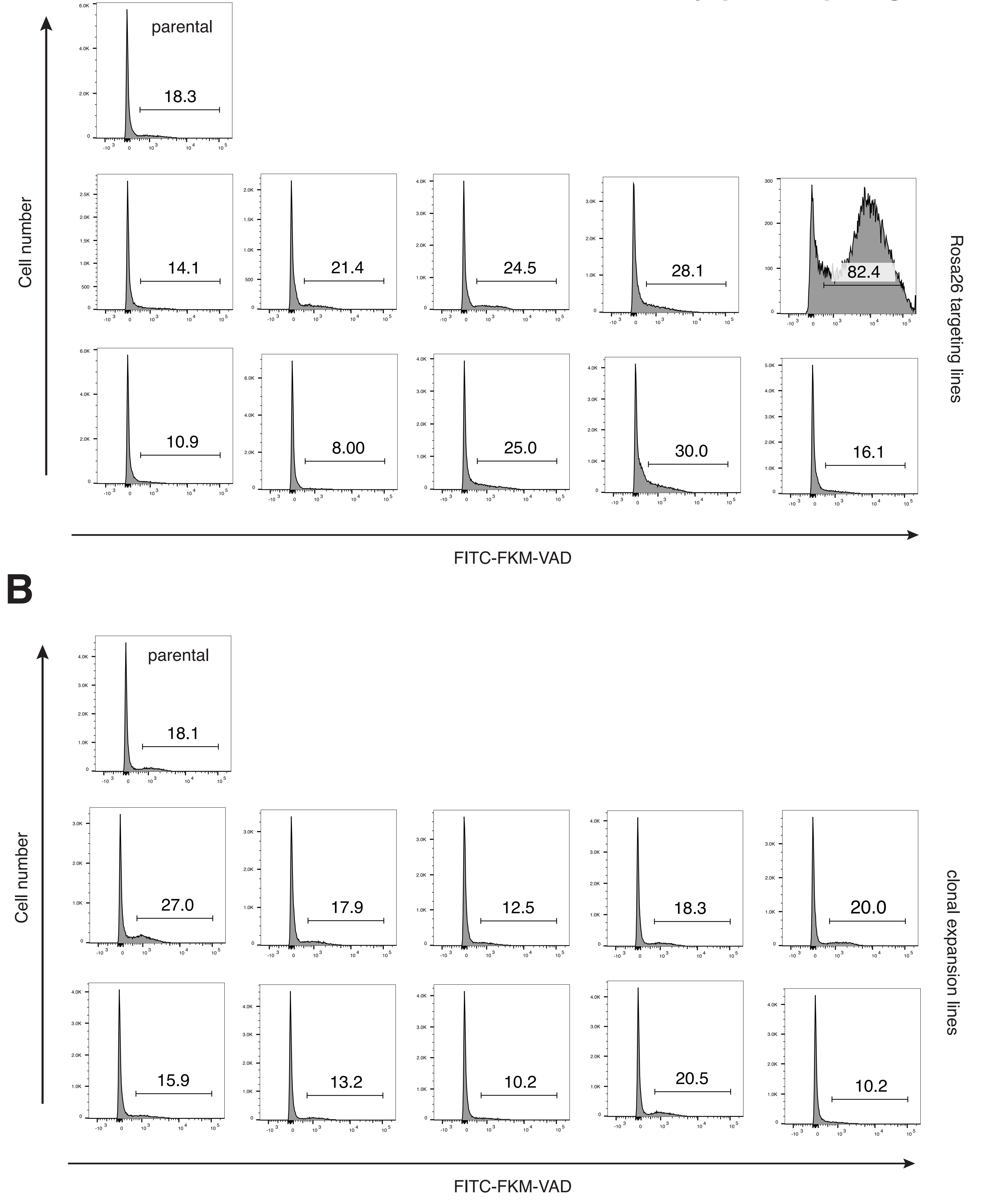
